# Neural correlates of consciousness in an auditory no-report fMRI study

**DOI:** 10.1101/2025.05.16.654468

**Authors:** Torge Dellert, Henning Balster, Insa Schlossmacher, Maximilian Bruchmann, Robert Moeck, Thomas Straube

## Abstract

In the search for neural correlates of consciousness (NCC), prominent theories disagree about the role of sensory versus wide-spread fronto-parietal brain activity. Research on auditory awareness has been widely neglected, and isolating NCC from correlates of task-related post-perceptual processes (e.g., report) is an ongoing challenge. The present study addressed these issues using functional magnetic resonance imaging (fMRI) during a no-report inattentional deafness paradigm. Sixty-three participants performed an auditory distractor task while supra-threshold but task-irrelevant sounds were presented in the background. Whereas one group was aware of these stimuli, another group remained unaware. Comparing brain responses to the critical sounds between aware and unaware participants while controlling for correlates of sensory and postperceptual processing revealed that auditory awareness was associated with significantly increased activity in secondary auditory but not in fronto-parietal areas. These findings suggest a dominant role of stimulus-specific sensory processing rather than widespread fronto-parietal information broadcasting in conscious perception.

## Introduction

Understanding how our brain gives rise to our experience of the world remains one of the biggest challenges for modern science. What are the neural correlates of consciousness (NCC)—the minimum neural mechanisms sufficient for any one specific conscious percept, for example, seeing an image or hearing a sound (Koch et al., 2016)? Recent research has focused on four prominent theoretical approaches (Seth & Bayne, 2022; Yaron et al., 2022), which strongly disagree about whether NCC can be localized in sensory or fronto-parietal brain areas (Cogitate Consortium et al., 2025; Mudrik et al., 2025). Most prominently, the global neuronal workspace theory (GNWT; Dehaene, 2014; Dehaene et al., 1998; Dehaene & Changeux, 2011; Mashour et al., 2020) argues that consciousness only occurs when perceptual information is broadcasted in a fronto-parietal network. A critical role of the prefrontal cortex (PFC; Panagiotaropoulos, 2024) is also proposed by higher-order theories (HOT; Brown et al., 2019; Lau & Rosenthal, 2011). In contrast, the recurrent processing theory (RPT; Lamme, 2003, 2006, 2010, 2018) argues that localized re-entrant activity within sensory cortices is sufficient to give rise to consciousness. This emphasis on sensory regions is shared by integrated information theory (IIT; Albantakis et al., 2023; Tononi et al., 2016).

Importantly, valid theories of consciousness should be generalizable across different sensory modalities. However, the vast majority of NCC studies has investigated conscious visual perception (Förster et al., 2020; Koch et al., 2016; Lepauvre & Melloni, 2021), while research on auditory perception is comparatively scarce (Dykstra et al., 2017; Snyder et al., 2015). Previous studies have linked auditory awareness to activity in auditory brain regions in Heschl’s gyrus and superior temporal gyrus/sulcus (STG/STS; Brancucci et al., 2011, 2016; Christison-Lagay et al., 2025; Dykstra et al., 2016; Giani et al., 2015; Gutschalk et al., 2008; Gutschalk & Dykstra, 2014; Königs & Gutschalk, 2012; Wiegand et al., 2018; Wiegand & Gutschalk, 2012) but also in fronto-parietal areas (Brancucci et al., 2011; Christison-Lagay et al., 2025; Eriksson et al., 2007; Giani et al., 2015; Sanchez et al., 2020; Wiegand et al., 2018) using functional magnetic resonance imaging (fMRI; Brancucci et al., 2016; Eriksson et al., 2007; Gutschalk & Dykstra, 2014; Wiegand et al., 2018; Wiegand & Gutschalk, 2012), magnetoencephalography (MEG; Brancucci et al., 2011; Giani et al., 2015; Gutschalk et al., 2008; Königs & Gutschalk, 2012; Sanchez et al., 2020; Wiegand & Gutschalk, 2012) and intracranial recordings (Christison-Lagay et al., 2025; Dykstra et al., 2016). In electroencephalographic (EEG) research, auditory awareness was associated with early negativities and late positivities in event-related potentials (ERPs; e.g., Eklund et al., 2019; Eklund & Wiens, 2019; Gregg & Snyder, 2012; Hillyard et al., 1971; Puschmann, Sandmann, et al., 2013).

However, most of these studies have systematically confounded conscious perception with task-related post-perceptual processing, such as decision-making, motor preparation, and report (Aru et al., 2012; de Graaf et al., 2012). Thus, it remains unclear which brain activity is necessarily required for auditory awareness per se (Dykstra et al., 2017). To tackle this problem, some visual NCC studies have employed no-report paradigms without trial-by-trial responses (Tsuchiya et al., 2015) and fMRI (e.g., Dellert et al., 2021; Frässle et al., 2014; Hatamimajoumerd et al., 2022; Kronemer et al., 2022). Previous studies addressing post-perceptual confounds in auditory perception, however, used EEG (Schlossmacher et al., 2021; Sergent et al., 2021; Zhu et al., 2024) with its limited spatial resolution, or did not contrast conscious and non-conscious processing of task-irrelevant stimuli (Wiegand et al., 2018), which is crucial to isolate NCC (Pitts et al., 2018).

Thus, the present no-report fMRI study aimed to identify neural correlates of conscious auditory perception with high spatial resolution and in the absence of decision-making. We manipulated awareness of task-irrelevant speech stimuli using inattentional deafness and analyzed brain responses during unaware and aware processing. Based on previous studies in other sensory modalities, we expected awareness-related activity in auditory but not fronto-parietal brain areas.

## Materials and Methods

### Participants

A sample of 63 right-handed participants (37 female, 26 male) was recruited from the local student community of the University of Münster via public advertisements. Their mean age was 23.35 years (*SD* = 2.94, range: 18–32). All had normal hearing and no psychiatric or neurological disorders. The sample size was chosen so that each group had approximately twice as many participants as previous fMRI studies on auditory awareness (Eriksson et al., 2007; Wiegand & Gutschalk, 2012). Participants provided written informed consent prior to the experiment and received monetary compensation. All procedures were approved by the local ethics committee. After exclusions (see Results), the final sample consisted of 55 participants (31 female, 24 male) with a mean age of 23.44 years (*SD* = 2.89, range: 19– 32).

### Apparatus and stimulus presentation

Stimulus presentation and response collection were controlled by Presentation (Neurobehavioral Systems, Inc., Berkeley, CA, USA). Auditory stimuli were presented using MRI-compatible insert earphones (HP AT01; MR confon GmbH, Magdeburg, Germany). The electro-dynamic headphone driver unit was placed next to the participants in the scanner bore and delivered the audio to their ears via air tubes and passive earbuds. Scanner noise was attenuated by hearing protectors worn over the earbuds. Visual information was projected onto a semitransparent screen positioned at the head end of the scanner. Participants viewed the screen in a mirror attached to the head coil and responded using a response box.

The stimulus presentation was physically identical throughout the experimental procedure (see below). At all times, a fixation cross in gray (RGB: 109/109/109) was presented on a black screen. The auditory stimuli consisted of three components we validated in a previous EEG study (Schlossmacher et al., 2021): beeps for the distractor task, critical speech stimuli for the awareness manipulation, and babble noise for masking. For an illustration of the waveforms, see Figure 1.

**Figure 1:**
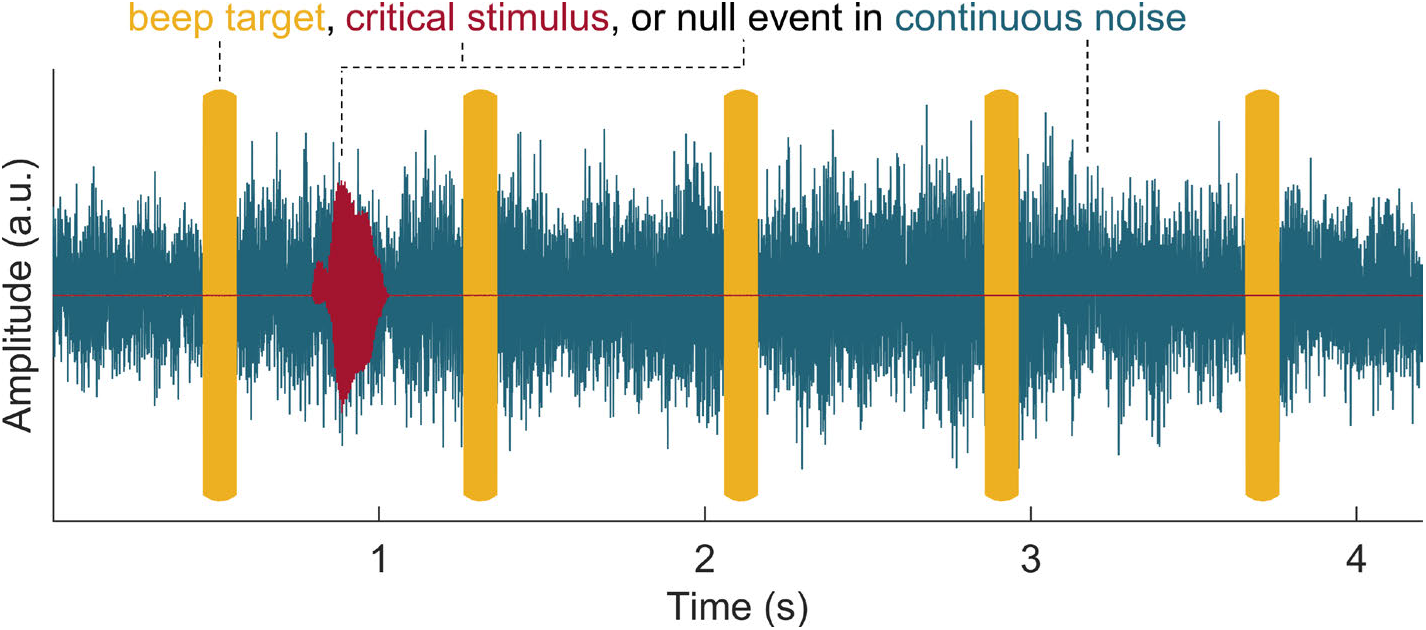
Stimulus presentation. Throughout the experiment, each trial contained continuous babble noise (blue) and beep tones (yellow). Moreover, each trial included one of the following three elements: an oddball beep target for the distractor task, a critical speech stimulus (red) with a jittered onset, or a matched null event. Due to the continuous presentation, the trial structure was unknown to the participants. During neuroimaging (phase 1), participants previously informed about the critical stimulus were aware of it, uninformed participants remained unaware of it, and both groups performed a distractor task detecting the beep targets. In a subsequent detection test (phase 2), both groups were asked to detect beep targets and critical stimuli in order to test their detectability.

Beeps had sinusoidal waveforms with a duration of 100 ms and frequencies of 800 Hz (standard) and 850 Hz (target), multiplied with a Gaussian waveform (duration = 100 ms, center = 50 ms, σ = 200 ms) to implement rise and fall times. The babble noise was taken from the Signal Processing Information Database (Johnson & Shami, 1993) and intensified by mixing the original noise with the same stream played backwards. The critical speech stimulus was the letter D (pronounced [deː] in German) spoken by a female voice, which was generated using a text-to-speech algorithm in MATLAB (Deng, 2020). The stimulus was cropped at the onset for optimal timing precision, resulting in a length of 230 ms, and a rise and fall time of 10 ms was implemented. The relative volume of the auditory stimuli was optimized in a pilot study, leading to an attenuation of the original beep, noise and critical stimuli to 76, 83, and 81 % of their original volume, respectively.

Beeps occurred every 800 ms, while the babble noise was presented continuously. Each trial was defined by the presence of either a beep target, a critical speech stimulus, or a null event. In beep-target trials, the first beep had a higher pitch (850 Hz) than the rest (800 Hz). However, participants were not aware of the trial structure due to the continuous presentation of beeps and noise. The onset of the critical stimulus randomly varied between 50 and 250 ms after the offset of the first beep so that it never overlapped with any beeps. The onsets of null events were drawn from the same random distribution, generating an adequate control event for each critical stimulus presentation. The order and timing of the trials was optimized for estimation efficiency in rapid event-related fMRI using optseq2 (Dale, 1999; Greve, 2009). For each experimental phase (see Procedure), ten different sequences were generated and randomly assigned to the participants. Overall, the inter-trial intervals (ITI) varied between 1.15 and 13.59 s (*M* = 3.61, *SD* = 2.32).

### Procedure

As described above, the stimulus presentation (Figure 1) was identical throughout the experimental procedure. However, participants were randomly assigned to two groups. While one group was informed about the presence of the task-irrelevant but critical speech stimuli (see below), the other group remained uninformed about them. This inattentional deafness procedure aimed to make informed participants aware and uninformed participants unaware of the critical stimuli during neuroimaging. Importantly, it offers increased experimental control and statistical power compared to previous inattentional blindness/deafness studies using a post-hoc classification of aware versus unaware participants (see also Pitts et al., 2018).

All participants were instructed to rest their gaze on the fixation cross, attend to the beep tones and to press the left button of the response box as fast as possible whenever they detected a deviant beep tone. Correct responses within 2 s were counted as hits. Moreover, participants were informed about the concurrent background noise being irrelevant and only there to increase the difficulty of the task. Before the experiment, participants completed three practice blocks, each including three target beeps but no critical stimuli. The beep-task difficulty was increased in each block by decreasing the target-to-standard frequency difference from 125 to 75 to the final 50 Hz. After each block, participants received visual performance feedback about the number of targets, hits, and false alarms.

Afterwards, the uninformed group directly proceeded with the neuroimaging phase (see below). The informed group, on the other hand, was shortly briefed about the occasional presentation of the spoken letter “D” during the upcoming experiment. In a short demonstration, they were presented with the babble noise and the critical stimulus accompanied by a visual presentation of the letter. It was emphasized that this stimulus was not relevant to the task, which only concerned the beep tones as practiced beforehand.

The subsequent neuroimaging phase lasted 15 minutes and included 100 critical sounds, 100 null events, and 50 beep targets. It comprised four blocks with mandatory breaks of at least 30 seconds after 50% and 15 seconds after 25% and 75% of the trials, respectively. During these breaks, participants were asked to relax their eyes and received visual performance feedback (the number of targets, hits, and false alarms) to maintain their motivation.

After neuroimaging (phase 1), we assessed participants’ awareness of the critical stimuli through their headphones. First, we openly asked them whether they had noticed any specific sounds in the background. If they accurately described the critical stimulus (i.e., “D” spoken by a female voice), they were classified as aware and asked during which block they had heard it first as well as how many times in total. If participants reported not to have noticed anything or only described sounds not matching the critical stimulus (e.g., scanner noise), they were classified as unaware.

After the awareness assessment, we implemented a detection test (phase 2) to ensure that all participants were generally able to perceive the beep targets as well as the critical sounds. To this goal, all participants were informed about the presence of both stimulus types in the upcoming task and asked to react to both of them as fast as possible using the left and right button, respectively. There was a short demonstration, during which the critical speech stimuli were accompanied by a visual presentation of the letter. Afterwards, participants completed a practice session including 5 beep targets and 5 critical stimuli, and received a visual performance feedback (targets, hits, and false alarms) concerning both the beep targets and the critical sounds. The main detection phase lasted 3.6 minutes and contained 24 critical sounds, 24 null events, and 12 beep targets. While it was designed for a behavioral test and not for fMRI analyses as it contained too few events, the same scanning sequence as in the neuroimaging phase was implemented to match the acoustic conditions including scanner noise. Performance feedback was displayed during a mandatory break of at least 30 seconds after half of the trials as well as at the end. Finally, participants were fully debriefed.

### Acquisition of neuroimaging data

MRI data were acquired using a 3-Tesla Siemens Magnetom Prisma with a 20-channel Siemens Head Matrix Coil (Siemens Medical Systems, Erlangen, Germany). First, a high-resolution T1-weighted scan with 192 slices was obtained for anatomical localization and coregistration (repetition time (TR) = 2130 ms, echo time (TE) = 2.28 ms, flip angle (FA) = 8°, field of view (FOV) = 256 × 256 mm, voxel size = 1 × 1 × 1 mm). A shimming field was applied in order to minimize magnetic field inhomogeneity. Subsequently, a functional data set consisting of 440 volumes with 42 slices was acquired using a T2*- weighted echoplanar sequence sensitive to blood oxygenation level-dependent (BOLD) contrast (TR = 2300 ms, TE = 30 ms, FA = 90°, FOV = 216 × 216 mm, voxel size = 3 × 3 × 3 mm).

### Analysis of behavioral data

Task performance was quantified by the sensitivity index d’ (Hautus et al., 2021) and response times (RTs) and analyzed in MATLAB. Due to the continuous stimulus presentation, false alarms were evaluated relative to the number of non-target time intervals of the same length as the 2-seconds response interval for hits (Bendixen & Andersen, 2013). To summarize single-trial RTs while protecting against the influence of skewness and outliers, the median was used (Wilcox & Rousselet, 2018).

For all further analyses, the following exclusion criteria were applied: Informed participants were excluded if they were unaware of the critical stimuli or if they only noticed them in the second half of phase 1 (neuroimaging). Uninformed participants dropped out if they spontaneously became aware of the stimuli during phase 1. Moreover, participants unable to detect the critical stimuli in phase 2 (d’ < 1) were excluded to ensure that the sounds were clearly audible for the final sample.

The detectability of the beep targets (phase 1 and 2) and, more importantly, the critical stimuli (phase 2) was analyzed by testing d’ against zero (one-tailed test) in each group. To investigate whether neural differences in the awareness contrast could also be attributed to interindividual differences in attentional allocation, d’ and RTs were additionally compared between aware and unaware participants (two-tailed test). T-Tests were computed using both frequentist and Bayesian inference (Keysers et al., 2020; Wagenmakers et al., 2018). Frequentist tests were regarded significant at *α* < 0.05. Bayesian tests used default priors (Morey & Rouder, 2011). Bayes factors (BF) quantify how much more likely one hypothesis is compared to another, with BF01 denoting the evidence for the null hypothesis and BF10 the evidence for the alternative hypothesis. We interpreted *BF* < 3 as anecdotal, *BF* ≥ 3 as moderate, *BF* ≥ 10 as strong, and *BF* ≥ 30 as very strong evidence (Jeffreys, 1961; Keysers et al., 2020). Effect sizes are reported as Cohen’s *d* (J. Cohen, 1988).

### Analysis of neuroimaging data

The MRI data were preprocessed using SPM12 (Wellcome Department of Cognitive Neurology, London, UK) and the Data Processing & Analysis of Brain Imaging (DPABI) toolbox (Yan et al., 2016) in MATLAB. The first five data volumes were removed to account for spin saturation effects. The remaining volumes were slice-time corrected and realigned using a six-parameter (rigid body) linear transformation. Anatomical and functional images were co-registered and segmented into gray matter, white matter and cerebrospinal fluid. Afterwards, the data were nonlinearly spatially normalized to Montreal Neurological Institute (MNI) standard space using DARTEL (Ashburner, 2007). Finally, they were resampled to 3 mm isotropic voxels and spatially smoothed using an 8 mm full width at half maximum Gaussian kernel.

For the first-level analysis, a general linear model (GLM) was estimated for each participant. A high-pass filter with a cutoff of 128 seconds was applied to remove slow signal drifts. Autocorrelations were modeled using SPM’s pre-whitening method FAST (Corbin et al., 2018) as recommended by Olszowy and colleagues (2019). The GLM design matrix contained the onsets of the critical sounds and null events. Predictors for standard and target beeps, responses and six head movement parameters were included as nuisance regressors. Onsets were convolved with a 2-gamma hemodynamic response function to model the BOLD signal change for each predictor. Contrast images (critical − null) of the beta estimates were created for each participant.

For the second-level analyses, we first investigated neural correlates of stimulus processing by testing the difference maps (critical − null) from the first-level analysis against zero in the unaware and aware group, respectively. More importantly, we isolated auditory NCC while controlling for sensory and postperceptual processing by comparing these difference maps between aware and unaware participants. For all tests, whole-brain analyses were performed using permutation tests with 10,000 sign-flipping permutations in the Permutation Analysis of Linear Models (PALM) toolbox (Winkler et al., 2014). We used threshold-free cluster enhancement (TFCE; Smith & Nichols, 2009), which includes spatial neighboring information without the need for setting an arbitrary initial cluster-forming threshold, and family-wise error (FWE) correction for multiple comparisons. Activations were considered significant if *p*FWE ≤ .05. In order to make potential sub-threshold activations transparent, the resulting statistical maps are presented using a dual-coded approach (Allen et al., 2012; Lieberman & Cunningham, 2009; Zandbelt, 2017), and non-significant activations with 0.05 ≤ *p*FWE ≤ 0.10 are reported additionally. Brain regions were identified based on the human Brainnetome Atlas (Fan et al., 2016).

## Results

### Behavior

After neuroimaging (phase 1), 27 out of 32 informed participants reported awareness of the critical stimulus. In contrast, all 31 uninformed participants remained unaware of it. The five informed but unaware participants were excluded as well as one who only noticed the critical stimuli in the fourth block. The remaining aware participants reported to have perceived the critical stimulus 58.23 times on average (*SD* = 35.18), but these estimations should be interpreted with caution. Two uninformed participants were excluded because they were unable to detect the critical stimuli in phase 2. Together, this resulted in samples of 26 informed, aware participants and 29 uninformed, unaware participants for the final analyses. In the following, ‘aware’ und ‘unaware’ always refer to these groups.

Results concerning task performance (d’ and median RTs) are presented in Table 1. In phase 1, target beeps were successfully detected by unaware (*t*(28) = 42.258, *p* < 0.001, *BF10* = 6.459 × 10^23^, *d* = 7.635) and aware (*t*(25) = 33.176, *p* < 0.001, *BF10* = 1.685 × 10^19^, *d* = 6.309) participants. The groups did not significantly differ in d‘ (*t*(53) = -1.680, *p* = 0.099, *BF01* = 1.153, *d* = -0.447), but RTs were significantly slower in aware compared to unaware participants (*t*(53) = 2.074, *p* = 0.043, *BF10* = 1.572, *d* = 0.552).

**Table 1:** Task performance as a function of group.

| Phase | Detection task | Sensitivity ($d'$) | | Median RT (ms) | |
| --- | --- | --- | --- | --- | --- |
|  |  | unaware | aware | unaware | aware |
| 1: Neuroimaging | Beep targets | 4.626 (0.589) | 4.341 (0.667) | 438 (58) | 472 (62) |
| 2: Detection test | Beep targets | 3.520 (0.549) | 3.700 (0.597) | 534 (99) | 536 (55) |
|  | Critical stimuli | 3.179 (0.742) | 3.299 (0.882) | 625 (87) | 640 (69) |

Values indicate $M$ ( $SD$ ). Note that ‘unaware’ and ‘aware’ always refer to the critical stimulus in phase 1 (neuroimaging). All participants in the final sample were aware of and able to detect the distractor beep targets in phase 1 and 2 and, more importantly, the critical stimuli in phase 2.

In the detection test in phase 2, formerly unaware (*t*(28) = 23.080, *p* < 0.001, *BF10* = 8.616 × 10^16^, *d* = 4.170) and continuously aware (*t*(25) = 19.064, *p* < 0.001, *BF10* = 4.914 × 10^13^, *d* = 3.625) participants demonstrated that they were able to detect the critical stimuli, suggesting that the sounds were well above the sensory threshold. The groups did not differ in d‘ (*t*(53) = 0.549, *p* = 0.585, *BF01* = 3.240, *d* = 0.146) or RTs (*t*(53) = 0.736, *p* = 0.465, *BF01* = 2.933, *d* = 0.196). Moreover, formerly unaware (*t*(28) = 34.543, *p* < 0.001, *BF10* = 3.110 × 10^21^, *d* = 6.241) and continuously aware (*t*(25) = 31.613, *p* < 0.001, *BF10* = 5.442 × 10^18^, *d* = 6.012) participants still successfully detected the beep targets in the dual-task setting of phase 2, and the two groups did not differ in d‘ (*t*(53) = 1.163, *p* = 0.250, *BF01* = 2.098, *d* = 0.310) or RTs (*t*(53) = 0.073, *p* = 0.942, *BF01* = 3.665, *d* = 0.019).

### Neuroimaging

#### Unaware group

During unaware processing, significant stimulus-specific responses were limited to auditory brain areas in the left and right STG (including the planum temporale) and Heschl’s gyrus. Statistical maps and beta estimates (critical – null) are presented in Figure 2 and statistics for local effects in Table 2.

**Figure 2.**
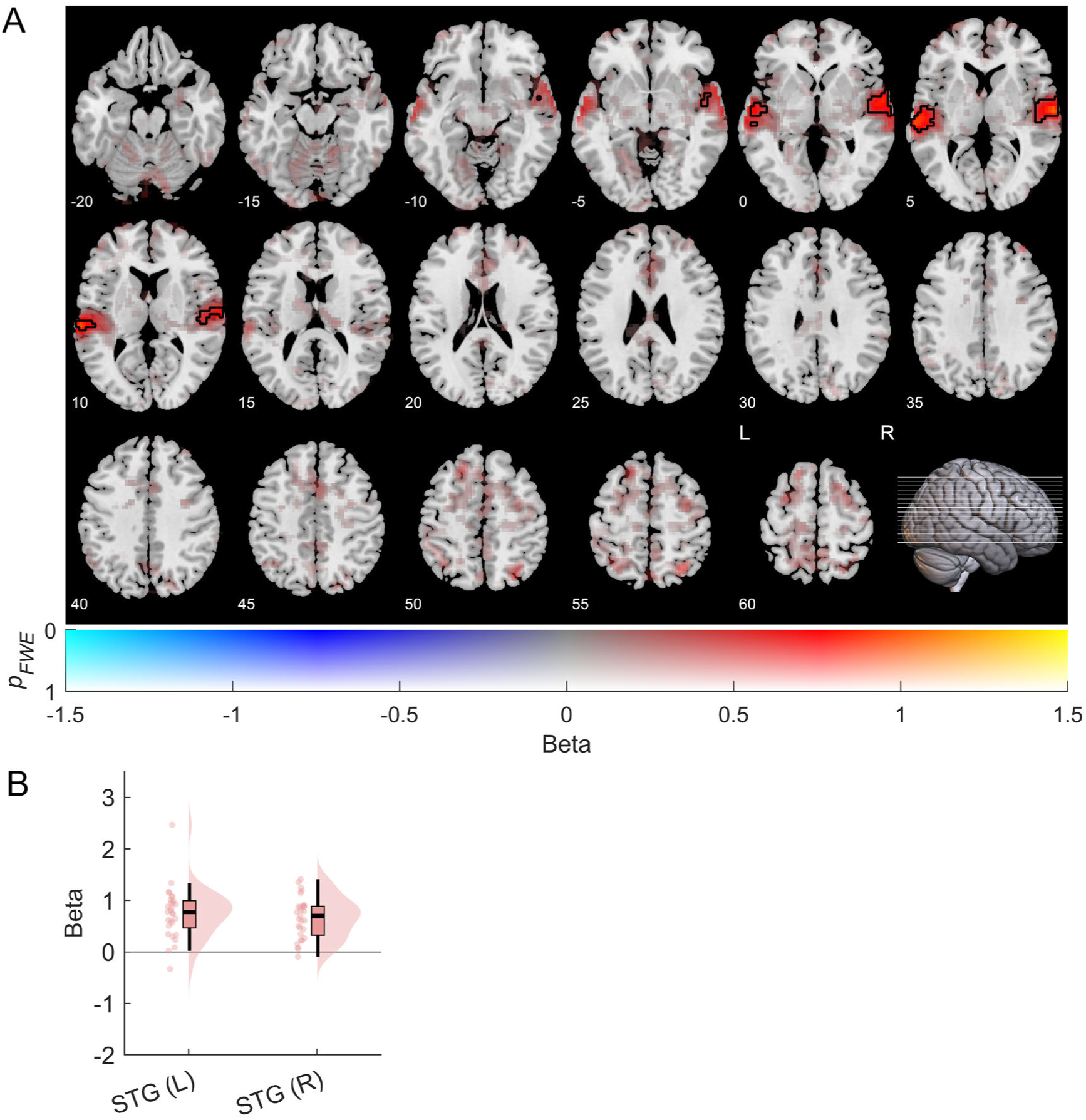
Neuroimaging results: Unaware group. Stimulus effects (critical – null, C – N) were tested against zero using permutation tests and TFCE. A) Statistical maps with beta differences (C – N) mapped to color hue and p-values (FWE-corrected) to transparency. Black contours mark activations with pFWE < 0.05. Axial slices are indicated by z coordinates (MNI) and lines in the sagittal view. B) Beta estimates (C − N) in peak voxels of significant effects (pFWE < 0.05) in the left (L) and right (R) superior temporal gyrus (STG) are presented using boxplots, half-violin plots, and scatter plots of individual data points.

**Table 2:**
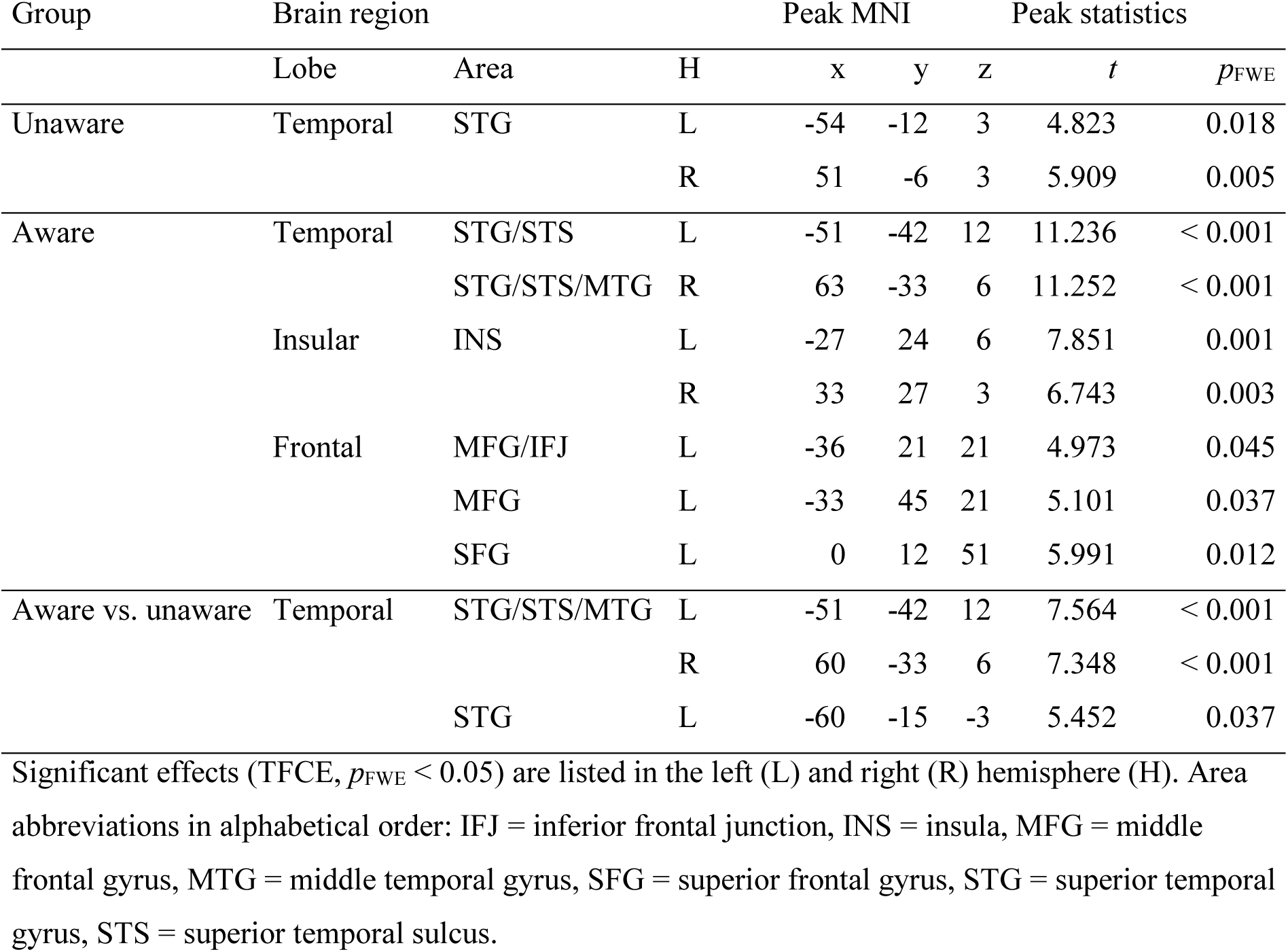
Neuroimaging results: peak statistics for whole-brain stimulus effects (critical – null, C − N) during unaware and aware processing as well as for awareness effects (aware (C − N) − unaware (C − N)).

#### Aware group

Aware stimulus processing also significantly activated auditory brain areas in the left and right STG and Heschl’s gyrus but additionally the bilateral STS and the left middle temporal gyrus. Moreover, significant effects were found in the bilateral insula and superior frontal gyrus as well as the left middle frontal gyrus and inferior frontal junction. Statistical maps and beta estimates (critical – null) are presented in Figure 3 and statistics for local effects in Table 2. Non-significant activations with 0.05 ≤ *p*FWE ≤ 0.10 during aware stimulus processing were observed in other parts of the inferior, middle and superior frontal gyrus, the inferior parietal lobe, and the precuneus.

**Figure 3.**
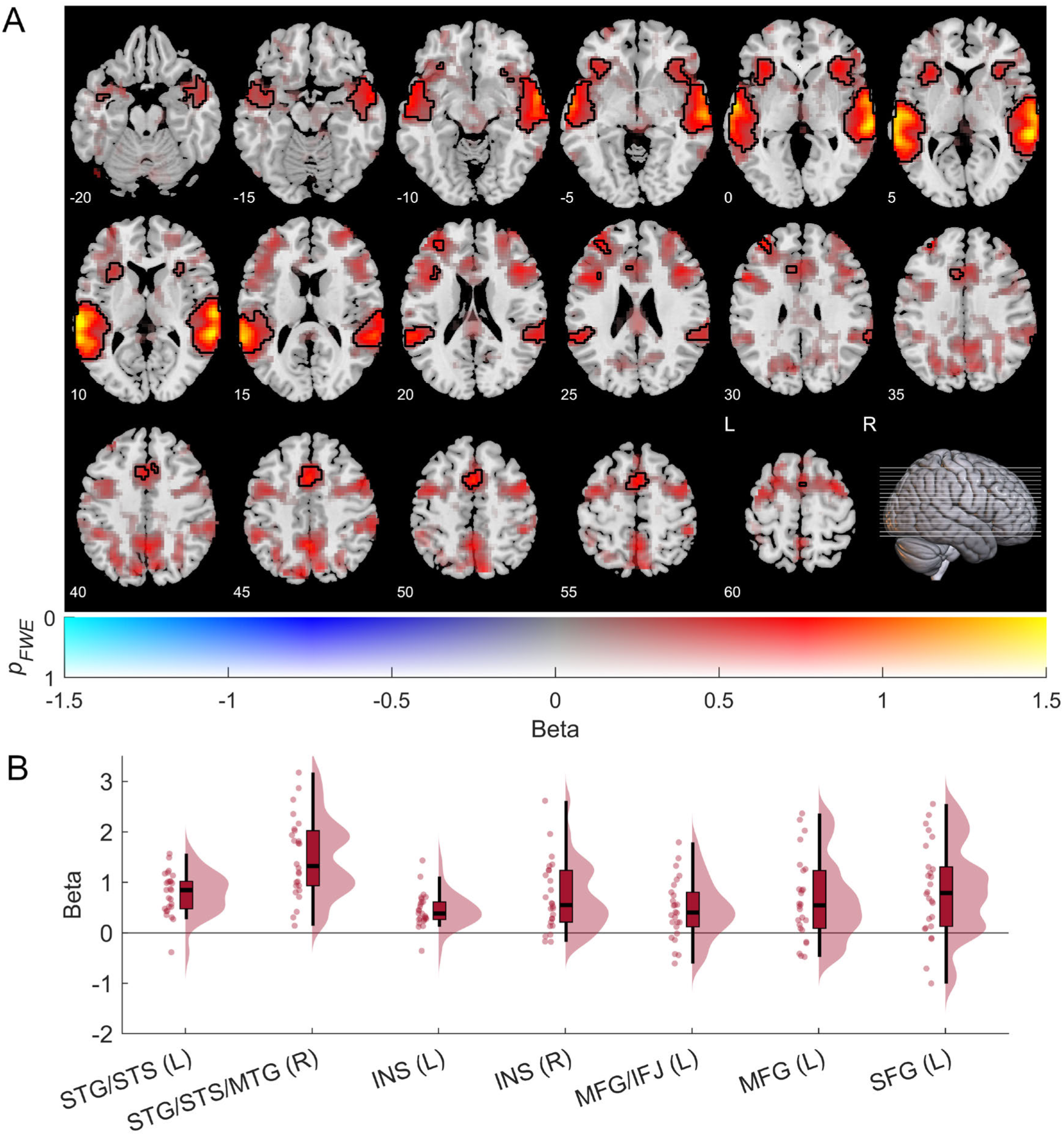
Neuroimaging results: aware group. Stimulus effects (critical – null, C – N) were tested against zero using permutation tests and TFCE. A) Statistical maps with beta differences (C – N) mapped to color hue and p-values (FWE-corrected) to transparency. Black contours mark activations with pFWE < 0.05. Axial slices are indicated by z coordinates (MNI) and lines in the sagittal view. B) Beta estimates (C − N) in peak voxels of significant effects (pFWE < 0.05) in the left (L) or right (R) hemisphere are presented using boxplots, half-violin plots, and scatter plots of individual data points. Area abbreviations in alphabetical order: IFJ = inferior frontal junction, INS = insula, MFG = middle frontal gyrus, MTG = middle temporal gyrus, SFG = superior frontal gyrus, STG = superior temporal gyrus, STS = superior temporal sulcus.

#### Aware versus unaware group

To isolate neural correlates of awareness, brain responses to the task-irrelevant critical stimuli versus null events were compared between aware and unaware participants. This contrast revealed that awareness was associated with significant, extensive increases of activity in the left and right STG, STS and anterior middle temporal gyrus. Statistical maps and beta estimates (critical – null) of awareness effects are presented in Figure 4 and statistics for local effects in Table 2. Additionally, non-significant awareness-related activity with 0.05 ≤ *p*FWE ≤ 0.10 was observed in parts of Heschl’s gyrus and the insula in the left hemisphere as well as in the bilateral inferior parietal lobule.

**Figure 4:**
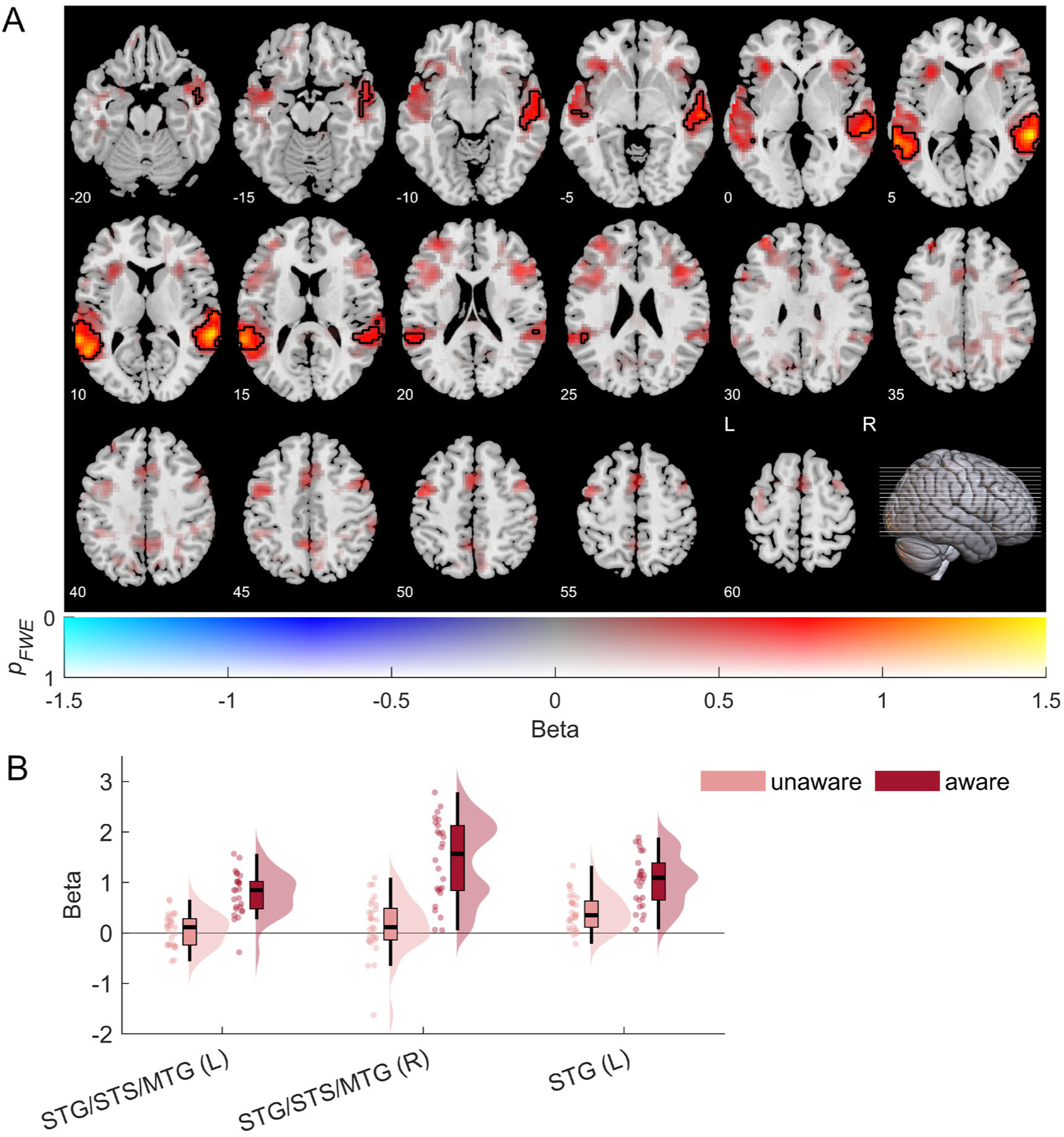
Neuroimaging results: awareness effects. Stimulus effects (critical – null, C – N) were compared between the unaware and aware group using permutation tests and TFCE. A) Statistical maps with beta differences (C – N) mapped to color hue and p-values (FWE-corrected) to transparency. Black contours mark activations with pFWE < 0.05. Axial slices are indicated by z coordinates (MNI) and lines in the sagittal view. B) Beta estimates (C − N) in peak voxels of significant effects (pFWE < 0.05) in the left (L) or right (R) hemisphere are presented using boxplots, half-violin plots, and scatter plots of individual data points. Area abbreviations in alphabetical order: MTG = middle temporal gyrus, STG = superior temporal gyrus, STS = superior temporal sulcus.

## Discussion

The present fMRI study aimed to isolate NCC in auditory perception in the absence of post-perceptual processes, such as decision-making and report. We manipulated awareness of task-irrelevant speech stimuli using inattentional deafness and compared stimulus-specific brain responses between aware and unaware participants, simultaneously controlling for effects of sensory and postperceptual processing. The results revealed that conscious auditory perception of task-irrelevant sounds was associated with strong activation of stimulus-specific auditory regions in the bilateral STG, STS, and middle temporal gyrus. In contrast, awareness-related activation in fronto-parietal regions was considerably weaker and non-significant, suggesting that a strong, widespread fronto-parietal ignition associated with global information broadcasting is unlikely to be a correlate of auditory consciousness. Using inattentional deafness (Dalton & Fraenkel, 2012; Macdonald & Lavie, 2011; Molloy et al., 2015; Raveh & Lavie, 2015) allowed us to manipulate awareness of physically identical, task-irrelevant sounds well above the sensory threshold by either informing participants about their presence in background noise or not. As a result, uninformed participants remained unaware of the critical stimuli.

After the awareness assessment, however, all included participants demonstrated their ability to detect the critical sounds.

Our fMRI results revealed that during unaware processing of the critical stimuli, activations were restricted to bilateral auditory regions including Heschl’s gyrus and the dorsal plane of the STG. These areas are sensitive to low-level processing of acoustic features (Alho et al., 2014; Binder et al., 2000; Okada et al., 2010), and the results agree with previous findings of auditory cortex activation during nonconscious processing (Bekinschtein et al., 2009; Boly et al., 2004; Colder & Tanenbaum, 1999; Laureys et al., 2000).

Aware processing of the critical stimuli activated the same core auditory regions but additionally the bilateral STS and the left middle temporal gyrus. These findings align with the role of these temporal regions in the perception of speech (Alho et al., 2014; Benson et al., 2001; Hickok & Poeppel, 2007, 2015; Jäncke et al., 2002; Vouloumanos et al., 2001), including whole sentences (Narain et al., 2003; Scott et al., 2000), words (Binder et al., 2000), syllables (Dehaene-Lambertz et al., 2005; Liebenthal et al., 2005), consonants (Obleser, Scott, et al., 2006), vowels (Obleser, Boecker, et al., 2006), and vocal sounds (Belin et al., 2000). Aware stimulus processing also recruited the ventral attention network (Fox et al., 2006; Vossel et al., 2014), which supports the detection of salient environmental changes (Kim, 2014). Specifically, significant effects were found in the bilateral insula (Menon & Uddin, 2010) and medial superior frontal gyrus (Yeo et al., 2011) as well as the left middle frontal gyrus (Vossel et al., 2014). Moreover, aware stimulus processing activated the left inferior frontal junction, an anterior core of the dorsal attention network associated with top-down attentional control (Kim, 2014). Taken together, the results accord with previous auditory fMRI studies (Alho et al., 2014; Hickok & Poeppel, 2007) even in the absence of a task (Block, 2019).

Importantly, to the present study’s main goal to isolate auditory NCC, whole-brain responses to the task-irrelevant critical stimuli relative to null events were directly compared between aware and unaware participants. This contrast simultaneously controlled for effects of sensory and postperceptual processing. Significant awareness-related activity was observed in secondary auditory regions (belt areas) in the bilateral STG, STS and middle temporal gyrus, which accords with previous studies on auditory awareness (Brancucci et al., 2011, 2016; Dykstra et al., 2017; Eriksson et al., 2007; Wiegand et al., 2018).

While prior research has linked auditory awareness to primary auditory cortex (Brancucci et al., 2011; Gutschalk & Dykstra, 2014; Wiegand & Gutschalk, 2012), we observed some activity both during aware and unaware stimulus processing, which suggests that it is not associated with awareness per se and highlights the importance of controlling for non-conscious processing. However, the discrepancy can also be explained by the content-specificity of NCC (Koch et al., 2016). While the abovementioned studies used low-level stimuli (i.e., tones), the present study featured speech stimuli known to be processed higher in the auditory hierarchy (Hickok & Poeppel, 2007).

In accordance with GNWT (Dehaene & Changeux, 2011; Mashour et al., 2020), previous research has also linked auditory awareness to frontal (Brancucci et al., 2011, 2016; Christison-Lagay et al., 2025; Dykstra et al., 2016; Eriksson et al., 2007) and parietal (Brancucci et al., 2011; Giani et al., 2015; Hill et al., 2011; Sanchez et al., 2020) brain activity. However, these studies could not separate awareness from task-related, postperceptual confounds (Aru et al., 2012; Snyder et al., 2015; Tsuchiya et al., 2015).

While one previous auditory fMRI study (Wiegand et al., 2018) associated fronto-parietal activity with task relevance, it did not allow to investigate NCC in the absence of it.

To our knowledge, the present no-report fMRI study is the first to isolate auditory NCC of task-irrelevant stimuli. Whereas the above-mentioned awareness effects in secondary auditory regions were strong and significant, any other activations including those in fronto-parietal areas of the dorsal and ventral attention networks (Kim, 2014; Vossel et al., 2014) were not. The strongest non-significant activation was observed in the left insula, which can be explained by its role in conscious auditory change perception independent of physical change (Puschmann, Weerda, et al., 2013) and in regulating brain network transitions gating conscious access (Huang et al., 2021; Kronemer et al., 2022). Together, these results do not support a key role of the lateral PFC in conscious perception (Brown et al., 2019; Mashour et al., 2020; Panagiotaropoulos, 2024).

While the left middle and superior frontal gyrus and inferior frontal junction showed significant stimulus effects in the aware but not the unaware group, the awareness contrast did not reveal significant effects in these areas. This was because sub-threshold frontal activity occurred even in the absence of awareness, cancelling according effects out in the NCC contrast. This finding highlights the importance of directly testing interaction effects in neuroscience (Nieuwenhuis et al., 2011).

In sum, our results support a dominant role of stimulus-specific, sensory rather than widespread, fronto-parietal activity in conscious perception in the absence of task relevance across sensory modalities (Dykstra et al., 2017; Gregg & Snyder, 2012). Thus, they are better compatible with sensory theories of consciousness, such as RPT (Lamme, 2003, 2006, 2010) and IIT (Albantakis et al., 2023; Tononi et al., 2016), than with cognitive theories, such as GNWT (Dehaene et al., 2006; Dehaene & Changeux, 2011; Mashour et al., 2020) and HOT (Brown et al., 2019; Lau & Rosenthal, 2011).

Notably, our findings agree with previous fMRI studies addressing post-perceptual confounds in auditory (Wiegand et al., 2018), visual (Dellert et al., 2021; Frässle et al., 2014; Hatamimajoumerd et al., 2022; but see Kronemer et al., 2022) and somatosensory (Peters et al., 2023; Schröder et al., 2019) awareness. Moreover, they are compatible with ERP studies (Dembski et al., 2021) linking early negativities over sensory areas to awareness and late centroparietal positivities to postperceptual processing in audition (Schlossmacher et al., 2021; Sergent et al., 2021; Zhu et al., 2024), vision (M. A. Cohen et al., 2020, 2024; Dellert et al., 2021, 2022; Pitts et al., 2012, 2014; Schlossmacher et al., 2020; Shafto & Pitts, 2015), and somatosensation (Schröder et al., 2021).

It is important to consider, however, that even fMRI cannot provide decisive evidence against awareness-related fronto-parietal activity, and conclusions should be made with caution. For example, weak effects in fronto-parietal brain areas may be explained by their interindividual heterogeneity (Marek & Dosenbach, 2018). However, overcoming the limitations of weak near-threshold stimuli, we manipulated awareness of sounds well above the sensory threshold (Dykstra et al., 2017) and prevented the masking of potential sub-threshold brain activity using dual-coded, transparent statistical maps (Allen et al., 2012). Thus, while such weak threshold-dependent effects in fronto-parietal regions might be involved in the modulation of conscious perception, it is safe to say that there was no evidence for a strong, non-linear “ignition” of these areas to be required for it (Mashour et al., 2020).

Next to these advantages, it is also important to point out limitations. First, the delayed awareness reports necessary to avoid post-perceptual confounds are not as precise as single-trial reports. Being surprised by the assessment, aware participants could only estimate the number of critical stimuli they consciously perceived. Moreover, putatively unaware participants may also have noticed but forgotten the stimuli. While such inattentional amnesia could lead to overestimating neural correlates of non-conscious processing and thus underestimating NCC, inattentional deafness has previously been shown to reflect perceptual rather than memory deficits (Durantin et al., 2017; Raveh & Lavie, 2015). Another limitation is that even no-report paradigms cannot avoid spontaneous post-perceptual processes associated with stimulus awareness (Block, 2019). Future studies should thus aim to reduce stimulus-related cognition as much as possible.

In summary, the present study suggests a dominant role of stimulus-specific, sensory rather than widespread, fronto-parietal activity in auditory consciousness when stimuli are task-irrelevant. Thus, our results inform current debates regarding the extent of involvement of these areas in conscious perception and emphasize the importance of deconfounding NCC from correlates of task-related post-perceptual processing across sensory modalities.

## Acknowledgments

We thank Lisa Müller for her support in data acquisition.

